# The flaws of fitness functions in changing environments

**DOI:** 10.64898/2026.04.27.720981

**Authors:** Loke von Schmalensee, Claus Rueffler, Lesley T. Lancaster, Greta Bocedi, David Berger

## Abstract

When predicting species’ responses to changing environments, one can use mathematical functions that describe how individual fitness components depend on the environment, or a single “composite” function that directly links fitness to the environmental state. The former approach is a cornerstone of process-based modelling, but the latter remains standard for developing fundamental theory and making ecological predictions. Yet, fitness is not a single instantaneous trait, but an integrated outcome of multiple underlying processes accruing throughout an organism’s life. We show that by ignoring the distinct environmental dependence of the underlying processes, predictions from composite fitness functions become inherently flawed in variable environments. We explore the magnitude of this error by leveraging empirical thermal reaction norms for four important life-history processes in an insect pest, the seed beetle *Callosobruchus maculatus*. We parameterize two fitness functions: one explicitly modelling the temperature-dependence of the four life-history traits independently (the “ground truth”) and one composite function, which treats fitness as a single, instantaneous outcome of the environment. By combining these two functions with hourly temperature data, we projected demographic responses under different warming scenarios across 300 sites over three beetle population origins (California, USA; Yemen; Brazil). We show that the composite function over- or underestimates fitness depending on subtle climatic differences and whether fitness is assumed to accumulate additively or multiplicatively, highlighting the problems of applying composite fitness functions to variable conditions. We conclude that explicitly modeling trait-specific processes will become increasingly important for accurate eco-evolutionary forecasting under future environmental change.

## Introduction

Due to human activity, organisms across the world are experiencing environmental changes at unprecedented rates. Land use intensification is causing drastic habitat changes^1,2^, pollution is creating harmful environmental conditions^2,3^, and anthropogenic climate change is rapidly shifting temperature distributions toward new extremes^4,5^. Via direct effects on organism fitness, these changes have dire consequences for biodiversity and ecosystem function^2,4,6–9^. Being able to predict such effects is therefore fundamental to forecasting eco-evolutionary dynamics and the associated socio-economic impacts of environmental change^10,11^. Theoretical and empirical work aimed at predicting organismal responses under changing environments has heavily relied on “composite” fitness functions, which map fitness onto an environmental gradient^12–21^. These functions are convenient, because they link the environmental variable of interest directly to a single fitness outcome. However, as we outline below, such composite fitness functions are inherently flawed for predicting fitness under variable environments.

Regardless of which fitness metric is adopted (e.g., intrinsic growth rate, *r*, net reproductive output, *R*_0_, or relative fitness, *w*), fitness represents a combined outcome of various physiological and/or behavioral processes (e.g., development, foraging, growth, reproduction)^22–24^. Empirically, however, composite fitness functions are often constructed by fitting a single curve to “direct” fitness measurements across a series of static environmental conditions, like *r* over salinity^25^, *R*_0_ over food availability^26^, or *w* over temperature^27^. In static environments, these composite fitness functions naturally align with the combined contribution of the underlying components. But nature is rarely static, and this agreement breaks down when organisms experience substantial environmental variation within their lifespans.

As demonstrated by Kingsolver and colleagues (2011)^28^ and reiterated since^29–32^, the shortcomings of composite fitness functions become obvious when considering how an organism’s environmental sensitivity might shift throughout ontogeny. For example, an organism whose juvenile life stages thrive under spring and fall temperatures, but whose adults perform best in summer, will have higher fitness being born in spring than in summer, even if the average temperature experienced throughout the life cycle would be the same in the two scenarios. A composite fitness function derived from a series of constant conditions would fail to capture this dynamic, since it effectively collapses the seasonally distinct responses into a single curve (Fig. 1A).

**Figure 1.**
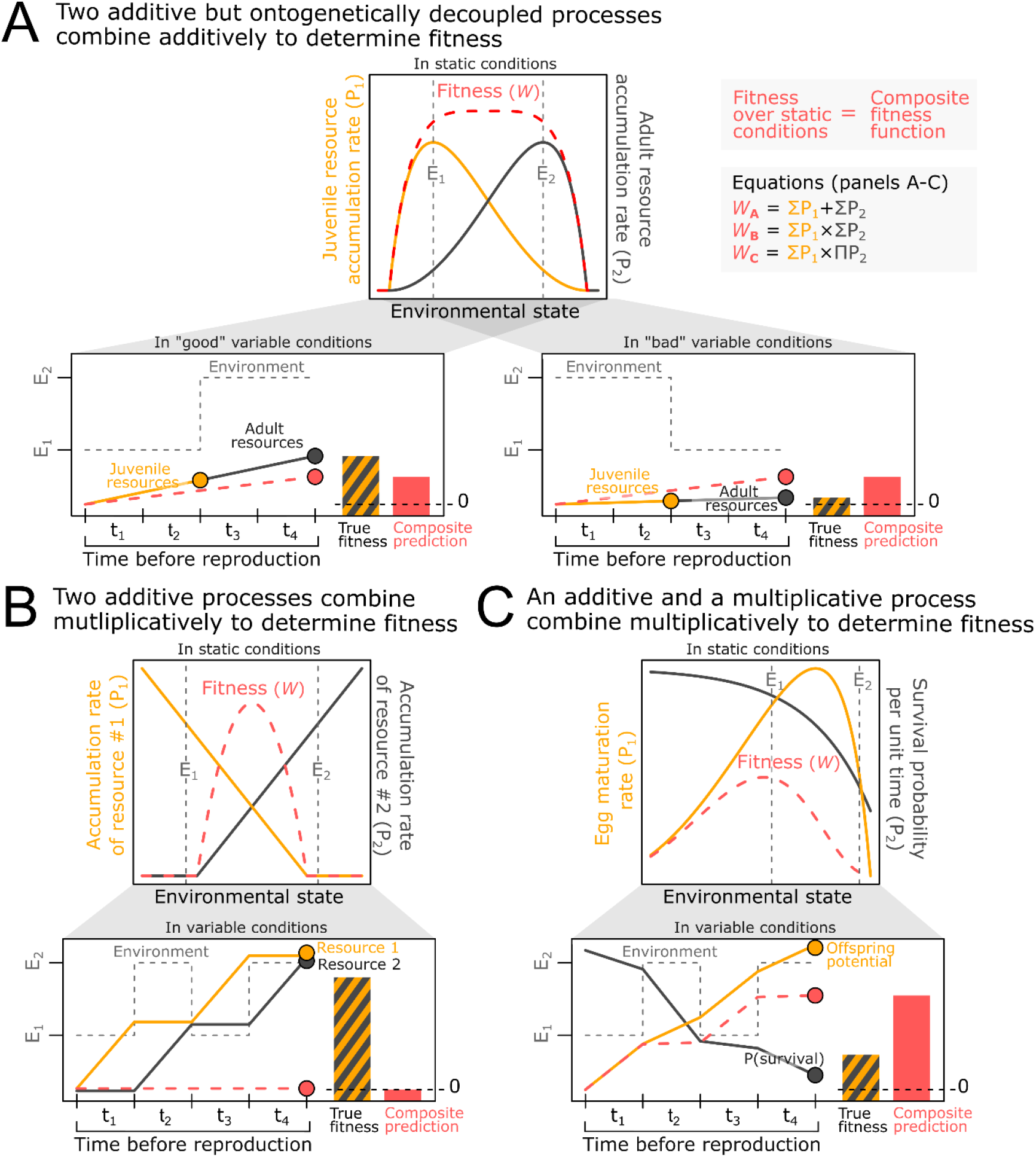
Examples of mismatches between true fitness and composite fitness functions under variable settings. Three hypothetical organisms with simple life-histories (black and yellow lines, top panels) highlight how composite fitness functions—representing fitness under static conditions (dashed pink lines, top panels)—fail to predict fitness under environmental variation (bottom panels). Note that the *y*-axes of the composite fitness functions in the top panels have been rescaled for clarity. The bottom panels illustrate how “fitness currencies” can accumulate before fitness is realized (t_1_–t_4_) in an environment alternating between two states (E_1_, E_2_). Bar charts compare true fitness, calculated by accumulating the separate component processes over time and combining them at the end of t_4_ (black/yellow lines and bars), against the prediction by the composite fitness function (pink bars and pink/dashed line). (A) Two processes contribute additively to fitness but are ontogenetically decoupled (juvenile and adult resource acquisition). Fitness is high only when environmental fluctuations align with the specific thermal requirements of the current life stage. In contrast, the composite function predicts identical fitness for both “good” and “bad” conditions since it ignores temporal sequence. (B) Two resources accumulate additively but determine fitness multiplicatively. In variable conditions, the organism achieves fitness by acquiring both resources sequentially. The composite function underestimates this by aggregating the instantaneously calculated combined fitness effect of the resources at each time step. (C) Fitness is determined by one additive process (offspring production) and one multiplicative process (survival). Here, the composite function overestimates fitness because it does not account for accumulated risk “leaking” into future conditions.

It remains underappreciated, however, that the inaccuracy of composite fitness functions under variable settings is not contingent on ontogenetic shifts in environmental sensitivity. This can be exemplified in ectothermic organisms, where external temperatures drive the rates of a multitude of physiological processes that jointly determine fitness^9,24,33,34^. These rate processes often progress over time towards a cumulative outcome—be it development rate determining the timing of life-stage transitions^35^, oviposition rates determining lifetime reproductive output^36^, heat injury accumulation rates determining survival probabilities_37_, or nitrogen fixation rates determining growth potential^38^. Because the interactions among these cumulative outcomes are not always instantaneous, composite fitness functions can substantially misrepresent fitness under environmental variation.

For instance, the utility of one nutrient may depend on the accumulation of another, driving down fitness in environments where either of the two is limited. In an environment that fluctuates between states each providing only one of two required nutrients, a composite fitness function will incorrectly predict zero fitness, as neither static state can indefinitely support life (Fig. 1B). A concrete, but less extreme, example of this phenomenon can be seen in the divergent thermal optima for photosynthesis and nitrogen fixation of some woody plants^38^. Similarly, mortality risk influences not only current but also future reproductive output. In a variable environment where high survival and high reproductive output coincide, fitness will be low for individuals that begin their life cycle in stressful conditions (e.g., a heatwave or a dangerous patch), as they may accumulate substantial mortality risk before realizing their reproductive potential. An instantaneous fitness function might correctly predict low fitness during the stress event but, as conditions improve, erroneously allow fitness to “catch up” by disregarding previous risk, leading to an overestimation of fitness (Fig. 1C). Finally, it is even possible for two organisms to exhibit the same fitness across all possible static conditions yet, due to underlying differences in individual fitness components, have different fitness under environmental variation (Fig. S1).

The above examples highlight the core problem: fitness functions parameterized across static conditions inherently assume that all fitness requirements are met simultaneously. Realized fitness, however, depends on the integration of separate processes over the organism’s entire life cycle. Here, we mathematically define fundamental mechanisms behind this discrepancy. We then present an empirical case study using thermal performance data from a cosmopolitan insect pest, the seed beetle *Callosobruchus maculatus*, to demonstrate how the use of composite fitness functions can lead to substantial prediction error depending on environmental context.

## Results

### Failure mechanisms of composite fitness functions

Decomposing fitness into constituent components makes it clear why composite fitness functions (e.g., functions derived from fitness measurements/calculations across static conditions) are unreliable for predicting fitness in variable environments. In Eq.1 and 2, we formalize this by contrasting the left-hand side, which represents the true fitness of an organism living through a variable environment from time 0 to T, against the right-hand side, which represents the mathematical equivalent of integrating a composite fitness function over that same period.

The most well-understood failure mechanism is “ontogenetic decoupling”, where the sensitivity of different life stages are decoupled so that the environmental influence on fitness varies with age (Eq. 1; Fig 1A) ^28–32^. In this case, fitness is maximized when favorable conditions for a given life stage co-occur with that specific life stage, whereas the same conditions may have negligible or even negative effects were they to occur at another time. In other words, under ontogenetic decoupling, the temporal covariance between environmental fluctuations and age-specific sensitivities plays a critical role in determining fitness.

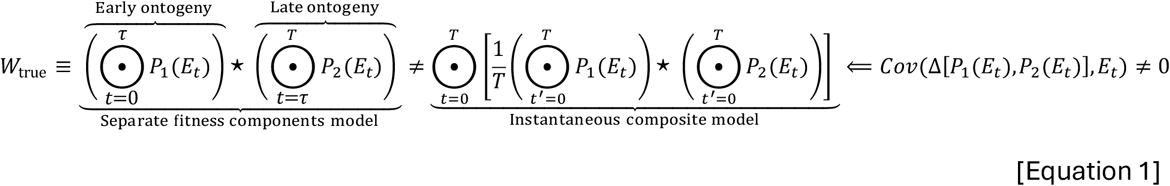

Here, we define true fitness (*W*_true_) as the result of two sequential processes (*P*_1_ and *P*_2_ ) that both depend on a time-varying environmental factor (*E*_*t*_). To generalize across different biological mechanisms, we use ⊙ as a general accumulation operator (e.g., summation for offspring production, or multiplication for survival) and ⋆ as a general combination operator describing how the cumulative outcome of *P*_1_ and *P*_2_ together determine fitness (e.g., additively for energy acquisition through two different processes, or multiplicatively for fertility probability and fecundity). On the left-hand side the “separate fitness components model” accurately calculates *W*_true_ by accounting for the temporal sequence of these processes. On the right-hand side, the “instantaneous composite model” evaluates fitness at each time point, *t*, by implicitly assuming that the current environment *E*_*t*_ persists over an entire lifetime, as represented by the dummy index *t*^′^. The resulting lifetime fitness values—each computed under chronic exposure to a single momentary temperature—are then averaged over the real time axis. This “instantaneous composite model” fails to represent *W*_true_ whenever the disparity between the processes (Δ) varies in response to a changing environmental state (*E*_*t*_) over the organism’s life. To maintain mathematical consistency, this disparity must be defined by the combination operator (⋆): if they combine additively, Δ represents the arithmetic difference (*P*_1_ − *P*_2_ ), whereas if they combine multiplicatively, Δ represents the ratio (*P*_1_ /*P*_2_ ). To exemplify: if survival in two successive juvenile life stages exhibits different thermal optima, the temperature sequence matters for an individual’s total survival. However, in the unlikely event that the two life stages retain a constant relative difference between their survival probabilities—meaning the thermal sensitivities are identical (note that the life stages can still have different *overall* survival)—the total survival probability will be independent of temperature sequence.

The failure of composite fitness functions is not solely dependent on age-structured responses. Even without ontogenetic shifts, an instantaneous composite function can misrepresent fitness under variable conditions. This occurs because the processes by which organisms accumulate “fitness currencies” (like growth, development, somatic damage, nutrients, or mating opportunities) and the processes by which these currencies interact to determine fitness are often not commutative (Eq. 2). That is, the result of combining the processes instantaneously and then accumulating this composite over time (as does a composite fitness function) is not necessarily equal to accumulating each process separately and combining them according to the specific structure the organism’s life-history dictates (Fig 1B-C). Put simply, the order of operations matters.

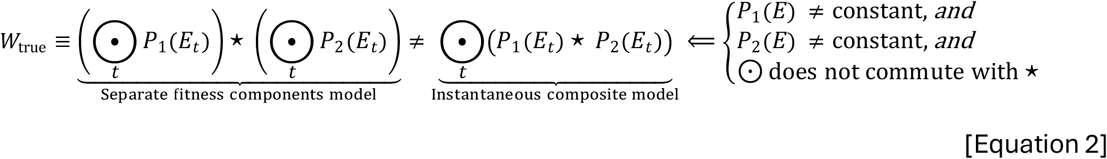

Here, the “separate fitness components model” on the left-hand side accumulates each process separately over time and then combines them, whereas the “instantaneous composite model” on the right-hand side combines the processes at each time point and then accumulates that composite. Thus, the “separate fitness components model” accurately represents true fitness (*W*_true_) by accounting for the compositional structure of the individual component processes. The “instantaneous composite model”, however, will only correctly calculate fitness if one of certain conditions are met, namely (1) all processes but one remain invariant over the environmental fluctuations or (2) all accumulation (⊙) and combination (⋆) processes are homomorphic (e.g., all are additive or all are multiplicative). Theoretically, the composite model also holds if the covariance between all component processes is zero over the range of environmental variation. However, this condition is near-impossible in practice and can effectively be ignored.

The environmental influence on an organism’s life-history is generally governed by a network of interacting processes, where some may be additive, others multiplicative, and where some might shift through ontogeny. Thus, Eq. 1 and Eq. 2 show that the general assumptions behind composite fitness functions are unrealistic under environmental variability. It should be noted, however, that the different failure mechanisms identified here may not all act in the same direction. As such, it is possible for the resulting errors to cancel each other out under certain conditions, creating an illusion of predictive accuracy. Therefore, a composite fitness function *could* generate relatively accurate estimates of fitness under certain scenarios, although this outcome cannot be expected to extend to all settings. Instead, adopting process-based models that explicitly account for the environmental sensitivities of the individual components, together with their compositional and temporal structure, provides a more robust approach.

### Thermal performance and fitness in seed beetles of three geographic origins under different climate warming scenarios

To explore the difference between composite and process-based approaches, we first assayed the thermal performance of laboratory populations of the agricultural pest beetle *Callosobruchus maculatus*, sourced from three geographic origins (California, USA; Brazil; Yemen) at five temperatures (17, 23, 29, 35, and 37°C). We recorded offspring development time, average offspring dry mass, and lifetime reproductive success of individual mating pairs (quantified by their production of offspring surviving to adulthood). We then fitted nonlinear temperature-dependent functions to the data using Bayesian methods (Fig. S2), which we converted to hourly rate functions of development, growth, survival, and fertility for subsequent integration across thermal regimes (Fig. 2A). See *Methods* for details.

**Figure 2.**
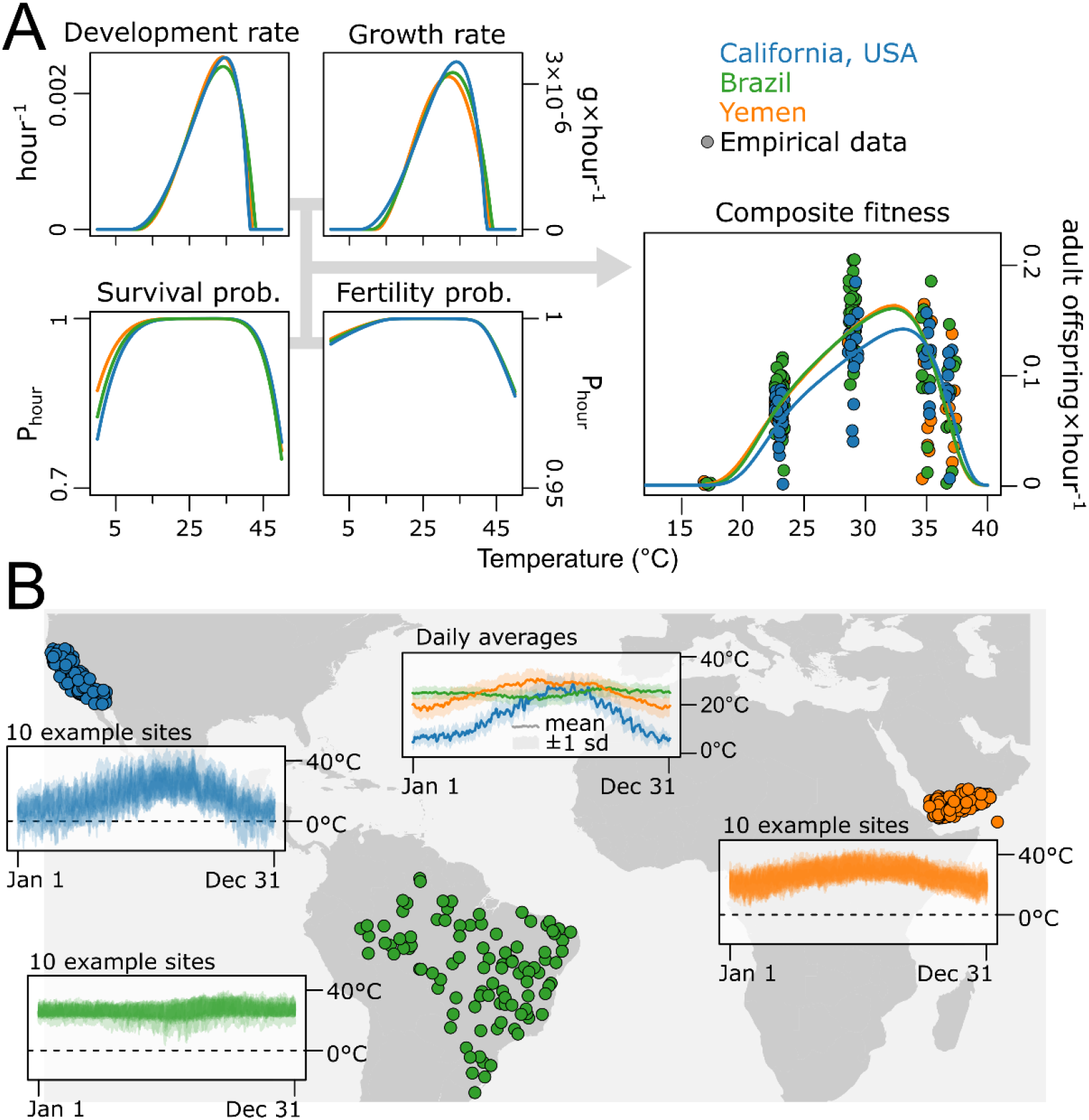
Empirical data from three geographic origins. (A) Temperature-dependent rate functions for *C. maculatus* populations sourced from California, USA (blue), Brazil (green), and Yemen (orange). Arrows illustrate how these components were combined to create the composite fitness function, here presented as an hourly rate of viable adult offspring production. Empirical data are shown for comparison, demonstrating the concordance between the separate component functions and their corresponding composite function under constant laboratory settings. (B) Map showing the 300 geographic sites (100 from each region) from which hourly ERA5-Land temperature time-series were sourced. Insets show 10 random hourly time-series from each region, and the central inset shows the mean annual cycle of daily average temperatures for each region, with shaded areas representing diurnal variation as one standard deviation around the daily mean.

For each origin, we parameterized a dynamic life-cycle model that integrates the separate hourly rate functions over time as a”ground truth”(hereafter the “separate-components model”). This separate-components model calculates fitness by dynamically simulating the full life cycle of an individual over a temperature time-series. Starting as an egg, the individual develops and grows according to its trait-specific reaction norms (Fig. 2A) driven by the sequence of temperatures it experiences (Fig. 2B). During development, the individual also accumulates potential risk of mortality and infertility from stressful temperatures. Once the individual reaches adulthood, its maximum fecundity potential is determined by its final size, and subsequent ambient temperatures dictate the hourly rate at which its egg pool is depleted. The average generation time (egg to egg) and the realized reproductive output (modulated by both maximal fecundity and mortality/infertility risk) combinedly determine fitness. From the separate trait-specific reaction norms, we also constructed origin-specific composite fitness functions (hereafter the “composite model”). These describe offspring production rate as a single, instantaneous function of temperature (Fig. 2A). Thus, in contrast to the separate-components model, the composite model fails to capture the temporal and compositional structure of the organisms’ life cycle. See *Methods* for details.

Lastly, to compare the two modelling approaches across a natural range of thermal conditions, we sourced hourly temperature time-series from 100 random grid cells (resolution: 0.1°*×*0.1°) within each geographic origin (300 grid cells in total) using the ERA5-Land dataset (Fig. 2B)^39^. To explore how the difference in predictions generated by the two modelling approaches could be affected by climate warming, we also simulated a range of warming scenarios for each unique time-series (present conditions up to +5 °C).

The three regions have distinct climate profiles: California shows high diurnal variability, seasonality, and heterogeneity, whereas Brazil is on average warmer and less variable; Yemen lies in between, with warm average temperatures but somewhat more pronounced variation than Brazil (Fig. 2B). We applied the composite and separate-components models for each genotype to the thermal regimes from their respective origin, calculating hourly finite growth rates^40^, *λ*_*hourly*_, over a full year. This metric represents generation-level fitness rescaled to a per-unit-time rate via an approximation of the Euler-Lotka equation^41^ (assuming density-independent growth and a stable age distribution, *Methods*). For each site, this yielded one prediction where fitness was determined instantaneously by the composite function, and a ground-truth estimation where fitness was calculated dynamically by simulating individual life cycles as previously described.

Comparing yearly fitness estimates from the composite and separate-components models revealed that the discrepancy between the two depends on both the thermal characteristics of the environment and the aggregation metric of choice—whether fitness is assumed to accrue additively or multiplicatively (Fig. 3). A typical “rate-summation” or “rate-averaging” approach^12–16,18,31,42–45^, which simply calculates the summed/averaged fitness over the year, yielded similar estimates for both models, except in sites with particularly unfavorable winters—mainly in California—where the composite model overestimated yearly fitness (Fig. 3A). Specifically, fitness was overestimated in sites where the separate-components model at some point reached a fitness of zero (Fig. S3). This is because the separate-components model dynamically integrates the individual processes over the full life cycle, increasing the risk of extinction during prolonged harsh periods (e.g., from the accumulation of thermal stress). In contrast, the composite model calculates fitness instantaneously, allowing it to accrue positive contributions during transient warm periods (e.g., warm winter days; see Fig. 4).

**Figure 3.**
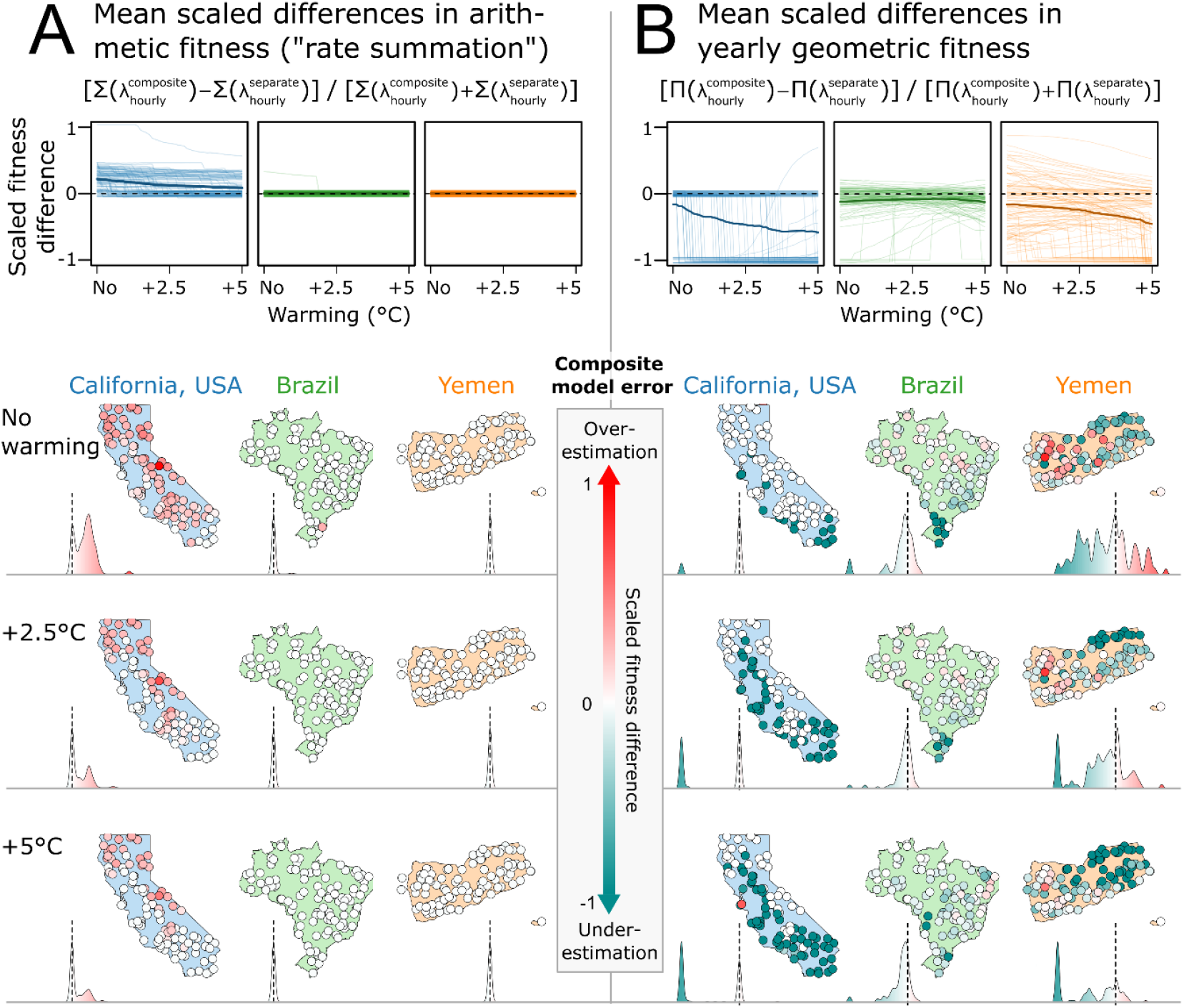
Differences between fitness estimates from the composite model and the separate-components model across thermal regimes. Scaled differences in *C. maculatus* fitness, calculated as (A) the hourly finite rates of increase summed over one year (“rate-summation”), or (B) geometric mean hourly finite rate of increase 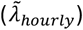 compounded over one year. Note that rate-summation disregards the compounding effects of population growth, and the geometric approach assumes density-independence; these metrics should thus not be interpreted as literal population projections. Values range from -1 (the composite model underestimates fitness, cyan) to +1 (the composite model overestimates fitness, red). Top panels show trajectories for individual sites (thin lines) and regional means (thick lines) across a 5°C warming gradient. Bottom panels visualize the spatial distribution of these differences at 0°C, +2.5°C, and +5°C warming for California (blue), Brazil (green), and Yemen (orange). Density plots below the maps show the frequency distribution of prediction errors for each region and scenario (dashed lines denote zero).

**Figure 4.**
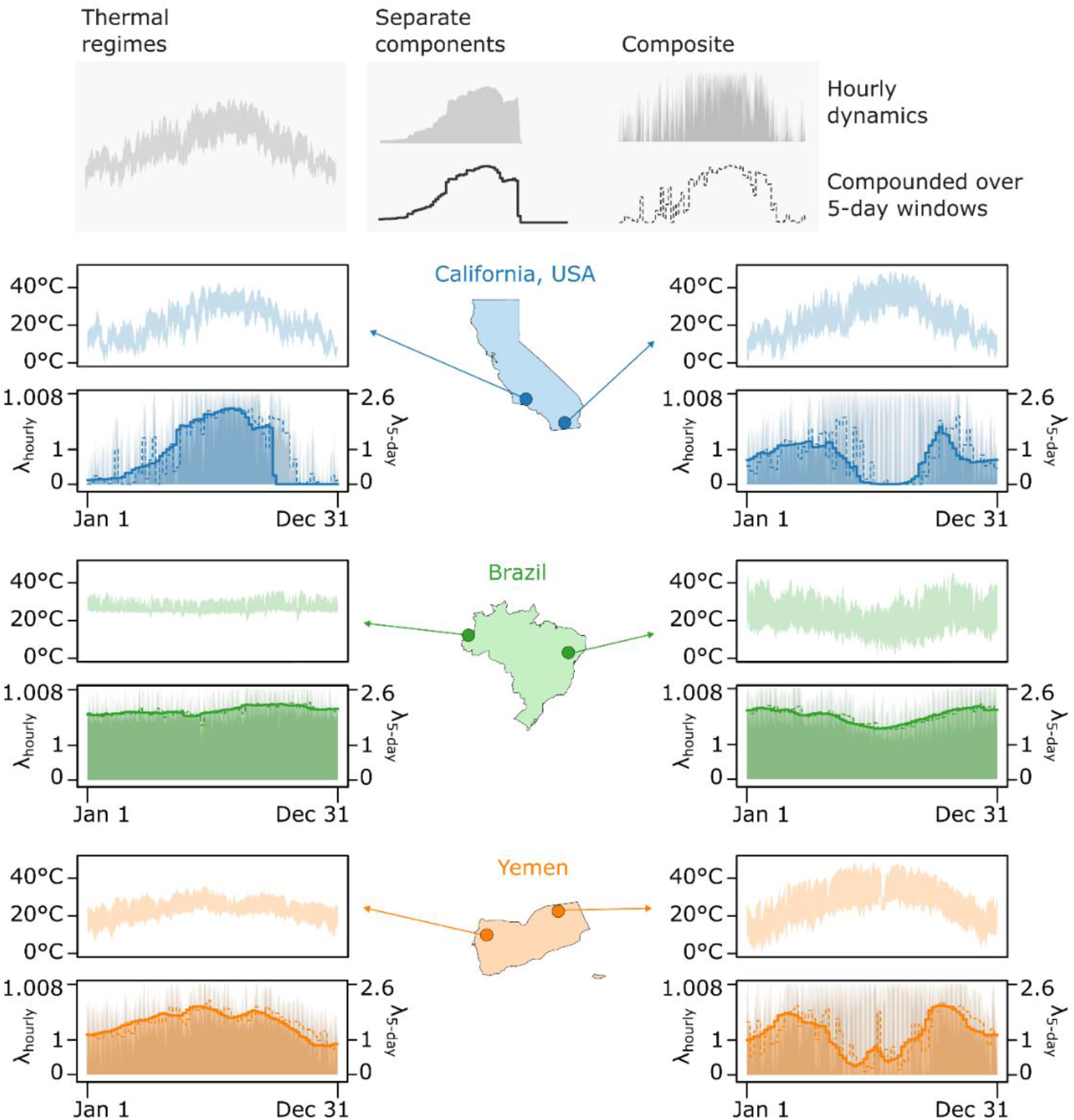
Seasonal fitness dynamics at example sites. Two example sites from each region—California (blue), Brazil (green), and Yemen (orange)—illustrate how discrepancies between the composite and separate-components models vary over a yearly cycle. For each region, top panels show temperature regimes (daily temperature ranges); bottom panels show the seasonal dynamics of the hourly finite rate of increase (*λ*_*hourly*_) predicted by the instantaneous composite function (dark shading, jagged fluctuations) and the dynamic separate-components model (light shading, smooth fluctuations). Lines show the product of *λ*_*hourly*_over non-overlapping 5-day windows (*λ*_5−*day*_), where solid lines represent the separate-components model and dashed lines represent the composite model.

When the annual fitness metric was calculated multiplicatively (instead of additively), the pattern changed (Fig. 3B). The same sites where differences in rate-summed fitness were substantial showed no discrepancies in geometric fitness between the composite and separate-components model (cf. Fig 3A & B). This is because the composite function inherently assumes a chronic exposure to each momentary temperature. Thus, the most extreme temperatures, which contribute to the cumulative failure in the separate-components model, are effectively treated as zero-fitness environments by the composite model. Consequently, in these specific sites the geometric fitness metric collapses to zero in both models—if a thermal regime leads to zero fitness over an organism’s lifespan (the separate-components model), if must necessarily include at least one single hourly temperature where *λ*_*hourly*_ would be zero also in the composite model. On the other hand, where the separate-components model predicts relatively low yet non-zero fitness—like southern California and north-eastern Yemen where summer temperatures get extreme (particularly with climate warming; Figs. 3,4)—the composite model often erroneously predicts extinction. This is because brief thermal extremes, which reduce but do not eliminate fitness in the separate-components model, are interpreted by the composite function as lethal chronic exposures, causing the geometric mean to crash to zero.

In sites with intermediate temperature variation (see Yemen in Figs. 2,4), the composite model’s error when predicting geometric fitness went in both directions (Fig. 3B). As the composite model responds instantaneously to short bursts of beneficial temperatures while simultaneously overestimating the impact of brief stress events (as seen in the temporal “jaggedness” of its predictions in Fig. 4) complex patterns of prediction errors arise depending on the frequency and amplitude of the fluctuations (Fig. S4).

To gain a better understanding of the dynamics behind the composite model’s estimation errors, we exposed the two prediction models to simulated temperatures with different diurnal and seasonal fluctuation amplitudes over a full year (Fig. 5). As opposed to the typical approach of fitting a curve directly to fitness measurements, we constructed our composite model bottom-up from the separate fitness components. While the resulting functions are the same (see Fig. 1A), our approach provides the unique opportunity to dissect the composite model and ascribe its prediction error to its four separate component traits, which revealed complex nonlinear effects of fluctuation amplitude and frequency. For arithmetic mean generation time and maximal fecundity, the composite model generally yielded estimates contributing to reduced fitness compared to those from the separate-components model. That is, the composite model overestimated generation time and underestimated maximal fecundity (driven by lower adult size estimates). However, this was not always the case under low to intermediate total temperature variability, where the pattern was sometimes reversed. In contrast, the composite model generally overestimated arithmetic mean lifetime survival and fertility compared to the separate-components model, but again, this pattern shifted at low to intermediate temperature variability around mean temperatures of 20°C and above.

**Figure 5.**
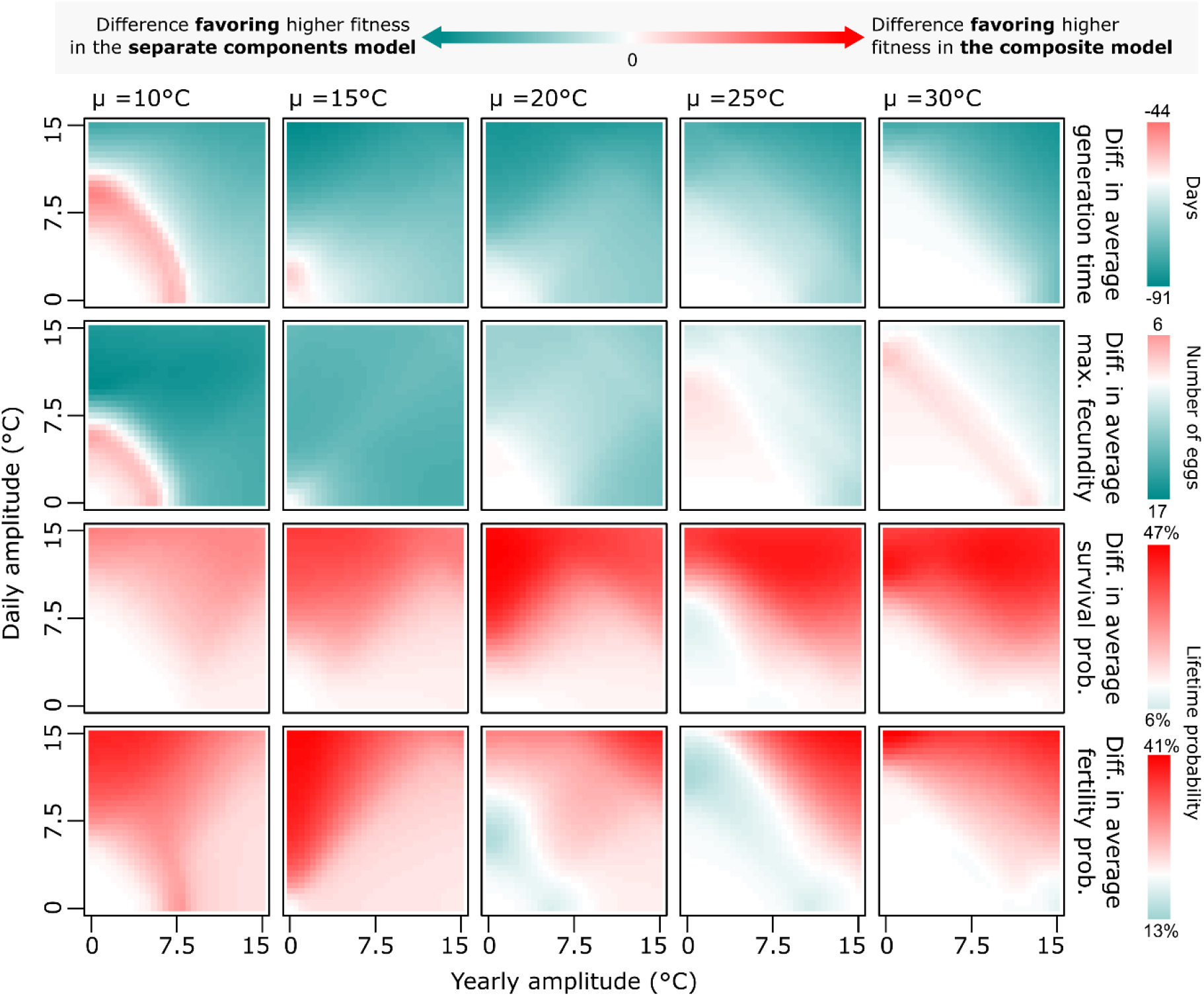
Trait-specific differences between the composite and separate-components model across simulated thermal regimes. Heatmaps show the difference in (arithmetic) yearly mean generation time (number of days; top row), maximum fecundity (number of eggs; second row), survival probability until adult maturation (third row), and probability of remaining fertile at adult maturation (fourth row). Each panel shows the interactive effects of yearly (x-axis) and daily (y-axis) temperature variation amplitudes for a given mean temperature (columns). White pixels represent equal trait-level predictions for both models. Red pixels constitute environments where trait values estimated by the composite model promote an overestimation of fitness (i.e., shorter generation time, or higher fecundity, survival, or fertility); cyan pixels signify an underestimation. Color bars show the minimum and maximum trait difference across all temperature treatments for each row, and these limits are denoted numerically at the tail ends.

The overall estimation biases of the composite model for the additively and the multiplicatively accruing traits (cf. the proportion of red vs. cyan in the top and bottom two rows of Fig. 5) stems from the difference between mathematically averaging instantaneous lifetime outcomes versus dynamically accumulating sequential rates and probabilities. For instance, development progresses very slowly near its thermal minimum; if temperatures briefly approach this minimum, the composite model’s instantaneous estimate of development time will be extremely high, inflating the mean development time across the time series. In the separate-components model, development simply pauses until temperatures rise again, preventing such extreme estimates. Conversely, the separate-components model correctly captures that survival is a product of sequentially experienced temperatures; encountering high mortality at any point during development reduces the overall lifetime survival probability. The composite model, however, averages independent lifetime survival outcomes calculated from each hourly temperature. Under harsh conditions, this inflates arithmetic mean survival estimates, as brief moments of benign temperatures (that in reality do not mitigate the damage of adjacent thermal extremes) are treated as parallel and independent full life cycles.

Together, the patterns in Fig. 5 show how composite fitness functions can distort the estimates of individual traits in distinct ways, with complex error landscapes arising from nonlinearities in the separate temperature responses. By extension, the net discrepancy between the composite model and the separate-components model (i.e., the composite model’s prediction error) depends on how the different component processes interactively influence fitness under a given environmental setting (Fig. S4).

## Discussion

Composite fitness functions, which map environmental states directly to instantaneous fitness outcomes, are often treated as a gold-standard for understanding fitness in dynamic settings^12– 16,18,31,42–45^. Yet, our findings show that in a naturally variable world, this assumption is fundamentally flawed. This is because composite fitness functions treat fitness as an instantaneous outcome, ignoring how individual component processes interactively operate over an organism’s lifespan (Fig. 1). As discussed below, this discrepancy between instantaneous composite estimates and actual fitness under variable settings can be misleading in several ways.

When fitness was aggregated additively over time, the composite model tended to overestimate *C. maculatus* fitness compared to the separate-components model (Fig. 3A), since it allowed fitness to accrue even during short bursts of favorable conditions under otherwise harsh periods(Fig. 4). When fitness was instead aggregated multiplicatively, the composite model’s errors were more diverse (Fig. 3B). In intermediately variable environments, the composite fitness function sometimes gave higher and sometimes lower fitness estimates than the explicit integration of the separate fitness components. However, in contrast with the additive aggregation approach, fitness was generally underestimated by the composite model in environments with (for *C. maculatus*) extreme seasonal conditions. This was because the composite model’s instantaneous, more jagged, response to temperature (Fig. 4) increased the likelihood of predicted extinction events, which, when fitness accumulates geometrically, carry over to subsequent time points.

Recently, Robey and Vasseur (2026)^45^ formally demonstrated how such extinction risk can be modulated by the temporal autocorrelation of temperature, with slower fluctuations increasing the probability of long bouts of harmful conditions driving a population to irreversible decline. However, using a composite fitness function obscures the underlying mechanisms behind such patterns^46^. A decline in *r* could be driven by stalled development (longer generation times), increased mortality, reduced fertility/fecundity or a combination of the above, each mechanism with potentially different eco-evolutionary implications^47,48^. For instance, Johnson and colleagues (2023)^31^ found that using a composite model may underestimate the detrimental effects of climate warming because it fails to capture how temperature-induced developmental delays prolong exposure to acute thermal mortality. Similarly, while composite fitness functions are used both as conceptual models^12^ and empirical evidence^20^ for inferring evolutionary responses, process- and trait-based approaches might provide a more accurate picture^11^. This is obvious when considering that organisms exhibiting equal fitness under a range of static conditions might still differ in their trait-specific reaction norms^49^ (Fig. S1) and therefore experience both unequal fitness and divergent selection pressures in environments that fluctuate at intragenerational timescales.

But where do we draw the line between a “true” fitness component and a composite? Temperature, for instance, acts directly on fundamental physiological processes^50^like metabolism^51^, photosynthesis^52^, and RNA transcription/decay^53^, which cascade to processes like growth^54^, mortality^55^, and development^56^, ultimately dictating fitness^20^. The question can be tackled both theoretically and empirically. Theoretically, our findings show that any set of traits which combine instantaneously, without carry-over into the future, can be compressed into a single composite response for modelling purposes. However, assigning processes to this category is a matter of judgement, as the validity of the approach of instantaneously combining processes ultimately depends on the timescale over which they interact, relative to the frequency of environmental fluctuations. Direct interactions between molecules where at least one has a very short half-life^57^emerge as conceivable examples where composite functions could be applied without much loss of information. Indeed, an empirically useful fitness component is simply one that can be adequately predicted under dynamic settings. In insect systems, for example, substantial progress is being made in leveraging models derived from static temperatures to accurately predict a range of physiological processes under variable conditions^55,58–63^.

These considerations also apply to our separate-components model. It is unlikely that the four processes we model herein are appropriate to treat as the sole”true” fitness components. Thus, our model is not a literal ground truth, but a baseline against which errors introduced by collapsing separate processes into a single instantaneous function can be revealed. Ironically, the implicit stable age distribution assumption behind the Euler-Lotka equation represents the same kind of temporal collapse we criticize in composite fitness models, since the true age distribution might vary in dynamic settings. While assuming stable age distributions under dynamic thermal regimes remains standard practice in the field^45,46,64,65^, a fully resolved approach would involve stochastic individual-based simulations. However, such models introduce their own assumptions and are beyond the scope of this study; our findings focus on the errors arising from collapsing separate component processes, upstream of the demographic model. Moreover, our results show that the discrepancy between a composite and separate-components modelling approach depends on which method is used to aggregate fitness over time. Fitness accrues geometrically, and this can theoretically be accounted for by integrating intrinsic, rather than finite, growth rate (*r* vs. *λ*) functions over time (since log *λ* = *r*). Yet, in practice such functions are often prevented from reaching *r* < 0 in environments where the population would naturally be in decline^16,45,65–67^, putting the approach functionally closer to an additive aggregation of fitness.

It should be noted that there are, indeed, cases when composite fitness functions naturally constitute appropriate approximations, even under realistic settings. For instance, it is trivial that composite models can predict fitness in organisms that do not experience much variation within their lifespan^68^, like single-celled organisms with short generation times in slowly fluctuating aquatic environments^69^. However, it is also plausible that predictions work better in single-celled organisms^70^ because they are less complex, putting fitness closer in the hierarchy to its fundamental physiological drivers^71^. Yet, species responses to environmental change are often intricate and context-dependent^72–75^, and the importance of models that incorporate the environmental dependence of trait-specific processes^76,77^—generating mechanistically grounded predictions beyond the conditions under which they were calibrated—will likely increase with climatic volatility^78^.

Although the focus here is on temperature, our framework applies to any environmental variable that fluctuates over individual organisms’ lifespans, whether through temporal variation, movement across spatially heterogeneous environments, or both^79^. Wherever such variation exists, relying on fitness functions—which map environmental states directly to high-level biological outcomes—risks mischaracterizing demographic and evolutionary responses. Instead, thoughtful integration of the diverse, trait-specific processes that govern an organism’s life cycle will yield a more nuanced eco-evolutionary understanding, ultimately enhancing forecast accuracy in naturally variable environments.

## Methods

### Animal rearing

The three laboratory populations of *C. maculatus*, originally sampled from Brazil, Yemen, and California, USA, have been maintained at population sizes of 300-400 adults and reared on cowpeas (*Vigna unguiculata*) under 55% relative humidity and 29°C conditions at Uppsala University, Sweden. For more details on the rearing procedure, we refer to Burc et al^80^. and Rêgo et al.^81^.

### Thermal performance and fitness

We assayed beetle responses across five constant temperatures (17, 23, 29, 35, and 37°C; 55% relative humidity; Panasonic MLR-352H-PE cabinets). First, we isolated beetle mating pairs in 60 mm diameter petri dishes with *ad libitum* cowpeas and then immediately moved them to the temperature treatments where subsequent oviposition occurred. For each mating pair, we scored lifetime reproductive success (LRS) by counting the total number of F1 adult offspring emerging from the host beans. We recorded the timing of the first emerging offspring (monitored daily) to calculate development time (time between oviposition start and first emergence). We also recorded mass as average offspring dry mass (using an Ohaus Pioneer PX85 balance). We performed these measurements at all temperatures for all three beetle lines yielding a minimal sample size at the lowest level of replication of n = 8 parental pairs (n_mean_ = 14, n_tot_ = 282). In all pairs with zero offspring, we sexed the parents through dissection, and for subsequent analyses we discarded data where same sex individuals had accidentally been paired (these are not included in the reported sample sizes).

### Curve fitting

We fitted thermal performance curves (TPCs) to the phenotypic data using a Bayesian framework in Stan^82^ via the R^83^ package brms^84^.

It is reasonable to model LRS as a Poisson process, but we noticed temperature-dependent deviations from this assumption in the form of zero inflation, particularly at extreme temperatures. This is likely because reproductive success is first determined by whether or not the parental pair can generate viable offspring at that temperature (i.e., both parents must be fertile). If the pair is fertile, the number of offspring produced is determined by the female’s fecundity. Moreover, the residual variance was not entirely homoscedastic on the log-scale (Fig. S2A). To account for both zero-inflation and among-temperature differences in residual variation, we therefore modelled LRS using a zero-inflated negative binomial error distribution. The temperature-dependence of the zero-inflation probability was modelled using a logit-linked refactored second-degree polynomial, allowing the probability of a pair being infertile to increase at extreme temperatures (Eq. 3).

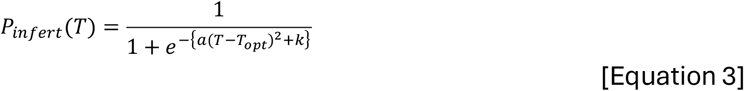

This function (Eq. 3) is defined by the parameters *a* (the curvature of the temperature-dependence of viability), *T*_*opt*_ (the temperature where viability is maximized), *k* (the minimal log-odds of infertility, at *T*_*opt*_ ), and the variable *T* (ambient temperature). The parameters were estimated jointly across geographic origins to aid model convergence.

To account for the unimodal and symmetrical relationship between LRS and temperature seen in our data (Fig. S2A), we constructed a custom nonlinear function modelling the temperature-dependence of fecundity, that is, the reproductive output per fertile beetle pair (Eq. 4).

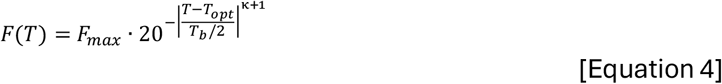

This temperature-dependent function *F* (Eq. 4) is defined by the parameters *F*_*max*_ (maximal number of offspring), *T*_*opt*_ (the temperature at which lifetime offspring production is maximized), *T*_*b*_ (the temperature range around *T*_*opt*_ where fecundity is > 5% of the maximum), κ (a parameter that regulates kurtosis, or the steepness of the decline in LRS away from *T*_*opt*_ ), and the variable *T* (ambient temperature). We estimated these parameters separately for the different geographic origins, except for the κ parameter, which we estimated jointly. The dispersion/shape parameter of the zero-inflated negative binomial distribution was estimated separately for each temperature.

For modelling the temperature-dependence of development rate (hour^-1^) and additive growth rate (g × hour^-1^), we employed the Lobry-Rosso-Flandrois function, which has a typical TPC shape and highly interpretable parameters (Eq. 5)^85,86^.

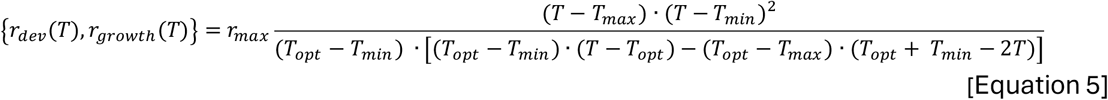

The temperature-dependent functions *r*_*dev*_ and *r*_*growth*_ (Eq. 5) are defined by the parameters *r*_*max*_(the maximal biological rate), *T*_*opt*_ (the temperature where *r*_*max*_ is achieved), *T*_*min*_ (the temperature below which the rate is estimated to be zero or negligible) and *T*_*max*_ (the temperature above which the rate is estimated to be zero or negligible), and the variable *T* (ambient temperature). To adhere to this logic, the mathematical function was modified using conditional statements to make it evaluate to zero at any input temperature below *T*_*min*_ or above *T*_*max*_ . We fitted the models using a lognormal error distribution and estimated separate parameters for each unique geographic origin.

Our models yielded good fits and posterior predictive checks revealed that the generative models could reliably approximate the distribution of the experimental data (Figs. S2, S6). When appropriate, we used informative priors to fit the models. For instance, *a*, the parameter describing the rate of change in infertility probability (away from the optimum) in Eq. 3, was bounded to be positive (as was the maximum development rate and fecundity). The temperature parameters were constrained to reasonable values using Gaussian priors. Additionally, to aid model convergence, the priors for the temperature parameters in the growth rate model were informed by the estimates from the development rate model(a closely linked trait). These priors were specified as Gaussian distributions with means corresponding to the posterior modes from the development rate model and a standard deviation of one. For all prior specifications, see Table S1.

### Derived functions

From these fitted curves, we derived functions describing the temperature-dependence of other important responses. Under constant temperatures these are defined as follows.

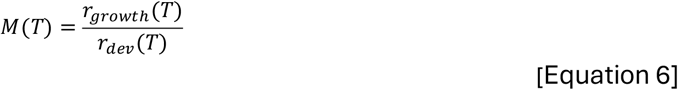

Function *M*(*T*) (Eq. 6) calculates the final adult mass by evaluating the static growth rate, *r*_*growth*_, over the required development time 1/*r*_*dev*_ at ambient temperature *T*.

Fecundity was maximized at 29°C and varied positively with adult body mass according to a power law (Fig. S7). We therefore estimated relationship of maximal potential fecundity and mass according to the function below.

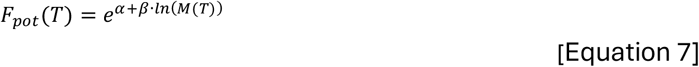

The function *F*_*pot*_ (Eq. 7) describes how the maximal potential fecundity of an individual depends on log-transformed body mass, *M*, where *α* is the intercept and *β* is the scaling exponent. We fitted this function using a linear model approach and removed non-significant terms (P > 0.05) for the final fit, leaving only *α* to be estimated separately for each geographic origin (Table S2).

Finally, to dynamically integrate our responses under variable thermal conditions, we converted them to hourly rates. For development rate and growth rate, no conversion was needed. We modelled oviposition rate based on a log-linear temperature-dependence found in previously published data (Eq. 8; Fig. S8)^80^.

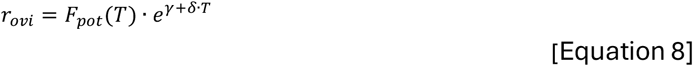

Here (Eq. 8), the hourly oviposition rate, *r*_*ovi*_, is a function of the maximal potential fecundity, *F*_*pot*_, and temperature, *T*, with an exponential intercept, *γ*, and slope, *δ*. We fitted this function using a traditional linear model approach and found no statistically significant effects of geographic origin (Fig. S8). Thus, parameters were estimated jointly across origins (Table S2).

We converted the probability of becoming infertile (based on egg count zero-inflation under chronic exposures, Eq. 3) to an hourly probability of remaining fertile, *s*_*fert*_, via the expected development rate.

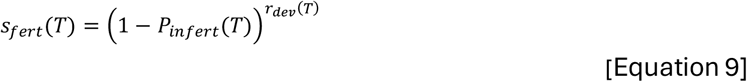

The function *s*_*fert*_ (Eq. 9) describes the hourly probability of maintaining reproductive viability at ambient temperature *T*. This transformation assumes that infertility risk accumulates proportionally throughout development, and scales the cumulative probability of fertility, 1 − *P*_*infert*_, by the hourly development rate, *r*_*dev*_, converting it to an hourly fertility probability, *s*_*fert*_ . To prevent mathematical artifacts where a development rate of zero would erroneously predict perfect fertility (since x ^0^ = 1), we constrained the development rate component; for temperatures outside the experimental range (<17°C or >37°C), the development rate was fixed to the values estimated at the boundaries.

Our modelling approach treats excess zeroes in the fecundity data (i.e., the portion of beetle pairs with zero offspring beyond what is expected from the negative binomial distribution) as infertility in the F0 mating pair. We attribute the remaining among-temperature variation in LRS to size differences in F0 individuals influencing fecundity potential (Eq. 7) and to F1 juvenile mortality driven by temperature stress according to Eq. 10.

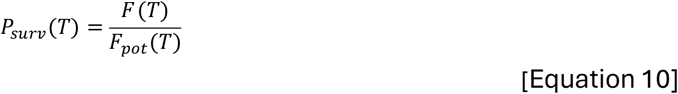

Here (Eq. 10), the total probability of surviving development, *P*_*surv*_, at ambient temperature *T* is simply expressed as the ratio of the modelled realized fecundity, *F*, to the maximal potential fecundity, *F*_*pot*_ . From this cumulative survival probability, we can then calculate hourly survival probabilities, *s*_*surv*_, following the same logic as for fertility rates (Eq. 11).

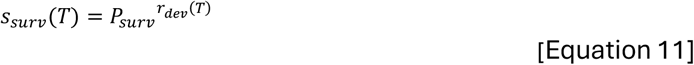

### Temperature data collection

To obtain realistic environmental temperatures for subsequent fitness estimations, we sourced hourly air temperature data from the ERA5-Land dataset provided by the European Centre for Medium-Range Weather Forecasts (ECMWF)^39^. We sampled time-series from 300 random geographic coordinates distributed evenly between the three geographical origins of our study populations: California, USA (n = 100), Brazil (n = 100), and Yemen (n = 100).

For each 0.1°*×*0.1° grid cell, we extracted a continuous two-year hourly temperature time-series. To ensure that our simulations were not biased by specific climate patterns in a single year, the two-year window for each time-series was randomized. This was achieved by randomly selecting a starting year between 2020 and 2023 for each location and extracting data for that and the following year.

Finally, to explore how differences in prediction between the two approaches were affected under different climate warming scenarios, we generated warmed thermal regimes by applying mean-shifts to the time-series at 0.1°C increments, from 0 to 5 °C.

### Approximating fitness via the instantaneous composite fitness function

In static and unconstrained (i.e., density-independent) settings, individual fitness can be estimated as the dominant eigenvalue (*λ*) of a Leslie matrix constructed from the reproductive schedule of just a single individual^40^. In other words, individual fitness can be described as the rate at which the individual would propagate copies of itself into the future, assuming all offspring inherit vital rates identical to those of their parents^87^. Because our temperature time-series have a 1h resolution, we use the aforementioned logic to estimate hourly individual growth rates, *λ*_*hourly*_, using an approximation of the Euler-Lotka equation (Eq. 12)^41^.

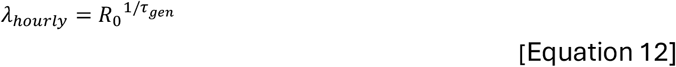

Here, τ_*gen*_ is the average generation time in hours (egg to egg; Eqs. 13-14), and *R*_0_ is the net reproductive output, that is, the number of viable female offspring per individual (Eq. 15). This approximation is fairly accurate since *C. maculatus* beetles spend most of their life in developing life stages, ovipositing for a fraction of their total lifetime. Under constant temperatures, development time, τ_*dev*_, is simply calculated as the inverse of the temperature-dependent development rate given in Eq. 5, *r*_*dev*_ (Eq. 13).

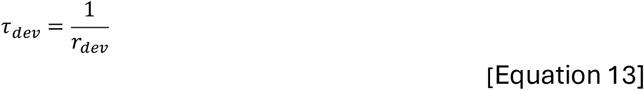

The remainder of the average generation time is defined by the mean timing of oviposition over the female’s reproductive lifespan. We assume beetles keep laying eggs with a temperature-dependent rate (Eq. 8) until their egg stores (*F*_*pot*_ ) are depleted. Because this rate is invariant under constant conditions, the average time of reproduction occurs halfway through the reproductive period. Thus, in static temperatures the mean generation time, τ_*gen*_, is given by the following linear equation (Eq. 14):

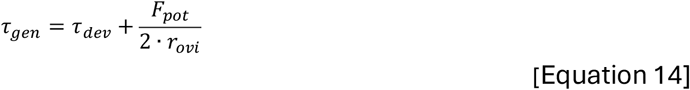

Since *C. maculatus* beetles lay eggs only for a limited period of their total lifespan, we assume that thermal stress accumulated throughout development determines the probability of becoming a reproductively viable adult, and that further effects of temperature stress are negligible once the adult stage is reached and adults can thermoregulate. Note that this simplification is supported by experimental data on fertility effects of adult and juvenile heat stress^88^. Thus, net reproduction (*R*_0_) is determined jointly by the thermal stress accumulated throughout development (*l*_*dev*_) and the reproductive potential (*F*_*pot*_ ), given by Eq. 15.

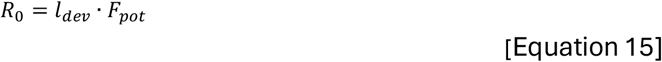

Here, *F*_*pot*_ is the temperature-dependent maximal potential fecundity given by Eq. 7, and *l*_*dev*_ is the temperature-dependent probability that an offspring is viable, that is, the parent beetle has remained both alive (*P*_*surv*_ ) and fertile (*P*_*fert*_ ) by the time of oviposition (Eq. 16).

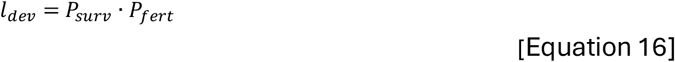

Under constant conditions, the probability of remaining fertile, *P*_*fert*_, is simply the complementary probability of becoming infertile given by Eq. 3 (1 − *P*_*infert*_ ). *P*_*surv*_ is given directly by Eq. 10.

### Approximating fitness via dynamic integration of fitness components

The formulations above provide a computationally efficient means to estimate fitness under constant thermal conditions and correspond to the “composite model” approach. However, natural thermal regimes are characterized by fluctuations that, due to nonlinearities in the temperature responses, preclude the use of static averages. To predict beetle fitness under these thermal regimes, we therefore developed a dynamic life-cycle simulation model based on numeric integration of the separate fitness components (the “separate-components model”). This model integrates our empirically derived hourly rate functions over the hourly temperature time-series to project the life cycle an individual egg initiated at hour zero of each day of the year. For each egg, we calculated expected outcomes of subsequent development, growth, fertility, mortality, and oviposition. Note that the life cycle of eggs laid on the last day of the year will spill over to the next year, hence the two-year temperature data.

Under variable temperatures, development time is dynamically determined by the continuous integration of the temperature-dependent development rate. To retrieve each individual’s specific thermal history, we therefore first calculated development time by integrating development rate until development was completed:

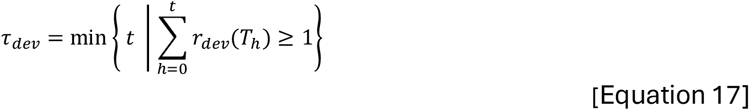

Here (Eq. 17), development time, τ_*dev*_, is measured as the minimum number of hours, *t*, required for the cumulative development rate ∑ *r*_*dev*_ to reach 1. This period, in turn, dictates the duration over which mass can accumulate (Eq. 18).

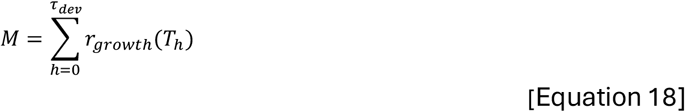

Again, we assume beetles keep laying eggs until their egg stores are depleted. Thus, the dynamic adult oviposition time is determined by the rate of oviposition, *r*_*ovi*_, and the maximal potential fecundity, *F*_*pot*_ (Eq. 19).

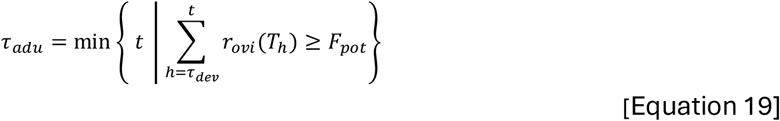

For each focal individual (i.e., each egg initiated on a separate day), we calculate an expected fertility probability, *P*_*fert*_, as a product of the hourly fertility rates, *r*_*fert*_, throughout development (Eq. 20).

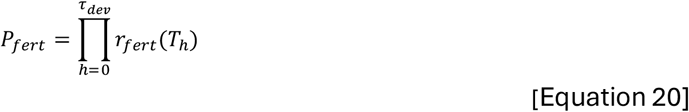

Similarly, we calculate expected individual survival probabilities, *P*_*surv*_, as a product of the hourly survival rates, *r*_*surv*_, throughout development (Eq. 21).

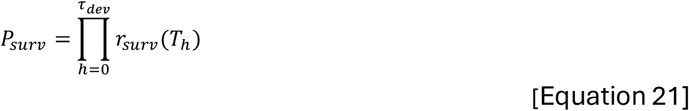

Finally, we calculate a weighted average generation time (τ_*gen*_) for the resulting offspring of each focal individual according to the formula below (Eq. 22).

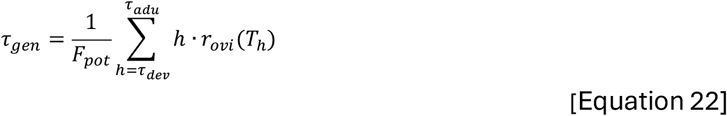

Here, *h* represents the age of the mother in hours, *r*_*ovi*_ (*T*_*h*_ ) denotes the number of eggs laid during hour *h* as a function of temperature, and the summation limits τ_*dev*_and τ_*adu*_represent the time of adult emergence and the time of egg pool depletion, respectively.

Having calculated the generation time (τ_*gen*_), maximal potential fecundity (*F*_*pot*_ ), and the cumulative probabilities of survival (*P*_*surv*_ ) and fertility (*P*_*fert*_ ), we estimated fitness, in terms of *λ*_*hourly*_, according to Eqs. 12, 15, and 16. Note that *λ*_*hourly*_ does not imply hourly population turnover; it is a rescaling of generation-level fitness to a per-unit-time metric, enabling direct comparisons and visualizations between the composite and separate-components models at the temporal resolution of the environmental data.

### Summarization

These approaches yielded two distinct time-series of theoretical hourly fitness values (*λ*_*hourly*_ ) for each thermal regime. To summarize the fitness over the full year period, we calculated both the yearly “rate-sums”, that is, *λ*_*hourly*_ summed over one year (representing the common approach of simply integrating/averaging a function over time^12–16,18,31,42–45^) and yearly finite rates of increase (*λ*_*yearly*_ ), that is, the product of *λ*_*hourly*_ over one year. While demographically inaccurate, the rate-sum (∑ *λ*_*hourly*_) serves as a proxy for “potential performance capacity” over a year, essentially treating fitness as a cumulative physiological trait. In contrast, the yearly finite rate of increase (*λ*_*yearly*_ ) provides a more realistic measure of potential population growth over a sequence of temperatures as it accounts for compounding effects. However, multiplying *λ*_*hourly*_ over a full year assumes unconstrained, density-independent growth throughout, which is unrealistic for natural populations. Both aggregation metrics should therefore be interpreted as indices of relative performance rather than literal demographic projections.

Standard relative error metrics, like percentage differences, are ambiguous when comparing finite rates of increase (*λ*) because their magnitude scales non-linearly depending on the time period over which *λ* is calculated. Therefore, to quantify the discrepancy between predictions from the composite and separate-components models, we instead calculated symmetric mean-scaled differences. For each thermal regime, we compared the two types of annualized outcomes predicted by the composite function (*W*_*composite*_ ) against those of the separate-components model (*W*_*separate*_ ) (Eq. 23).

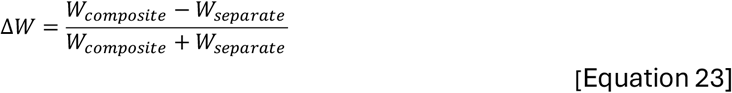

The resulting index is bounded between -1 and +1, providing a standardized error metric where zero indicates perfect agreement between the models, positive values indicate overestimation by the composite model and negative values indicate underestimation by the composite model.

### Analysis of life-history traits under simulated thermal regimes

Finally, to systematically explore how within- and between-generation thermal variability affects the discrepancies between composite and separate-components models, we conducted a sensitivity analysis using synthetic temperature regimes. We generated hourly temperature time-series from a mean temperature, *µ*, and cosine functions representing yearly and daily variation, with amplitude A_*yearly*_ and A_*daily*_, respectively (Eq. 23).

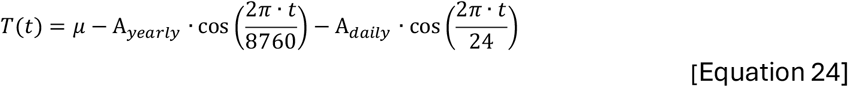

To illuminate underlying patterns behind the discrepancies between the composite and separate-components models, we isolated and compared their predictions for individual traits, averaged over a full year (arithmetic means). The separate traits were generation time (days), maximum fecundity (eggs), lifetime survival probability, and lifetime fertility probability. We then calculated the differences in these trait means between the two models and averaged the differences across the three geographical origins. Using this methodology, we constructed an array of error landscapes with mean temperatures ranging from 10°C to 30°C in 5°C increments, and yearly and daily fluctuation amplitudes ranging from 0°C to 15°C in 0.5°C increments.

