## Supporting information for "The flaws of fitness functions in changing environments"

### Contents

Figures S1–S8 (pp. 2–9)

Tables S1–S2 (pp. 10–11)

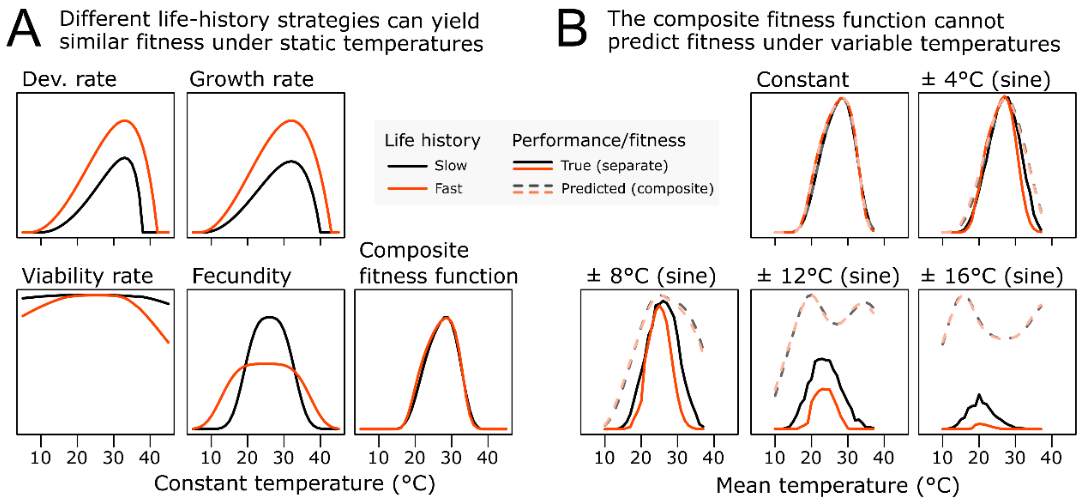


**Figure S1**. Thermal reaction norms for four interacting traits and their combined outcome in two hypothetical organisms with contrasting life-history strategies: a "fast" strategist with rapid growth and development (orange lines), and a "slow" strategist with slower growth but higher potential lifetime fecundity (black lines). Although the underlying processes differ in their reaction norms, they combine to produce nearly identical composite fitness functions under constant conditions. (D) Comparisons of true fitness (solid lines) versus predicted fitness from the composite fitness function in C (dashed lines) under sinusoidal temperature fluctuations. As thermal variability increases, the divergence between true and predicted fitness demonstrates the poor applicability of composite fitness functions under variable settings.

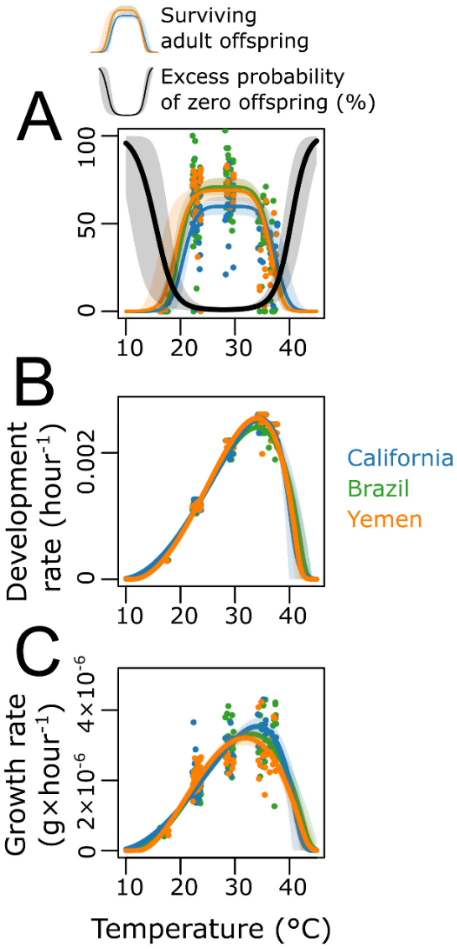


**Figure S2**. Fitted curves to the empirical data (points). Points and curves are colored by origin: California, USA (blue), Brazil (green), and Yemen (orange) (A) Plot showing the fit for the model of lifetime reproductive success. The black line represents the probability of infertility, that is, the excess probability of a beetle pair having zero viable offspring. The colored lines represent the fecundity, describing the expected number of offspring surviving to adulthood, under the condition of fertility. (B) The fitted curves describe how development rate varies with temperature. (C) The fitted curves describing how growth rate varies with temperature.

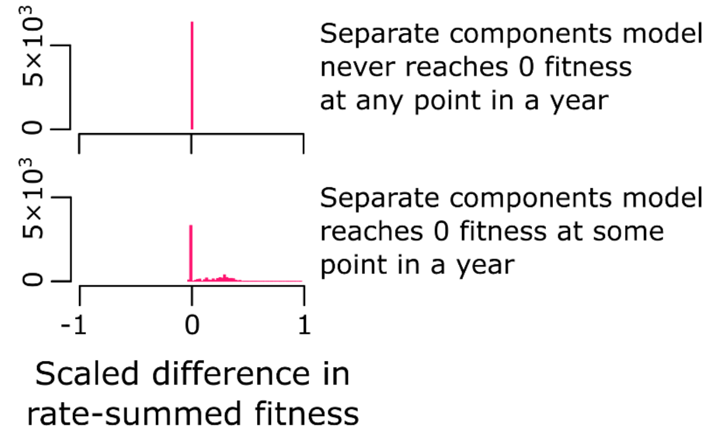


**Figure S3**. The scaled differences in arithmetic fitness (“rate summation”; values above 1 indicate an overestimation by the composite fitness function) between the composite and separate components models in sites where hourly fitness in the separate components model never reaches zero (top panel) or does so at some point (bottom panel). Note that differences in arithmetic fitness are always positive and only arise in sites where the separate components model predicts periods where fitness is zero (during which time the composite model keeps accumulating).

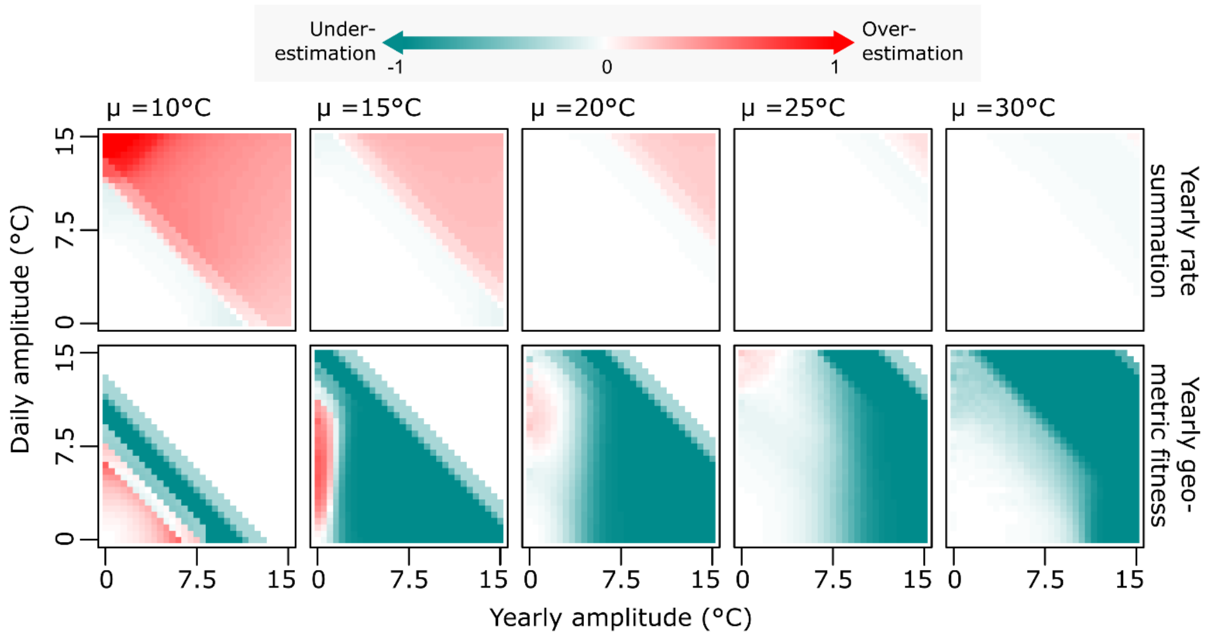


**Figure S4.** Scaled fitness differences under simulated thermal regimes. Heatmaps show the mean scaled fitness difference between the composite and separate-components models (as in main Fig. 3), across different mean temperatures (columns) and gradients of yearly (x-axis) and daily (y-axis) temperature variability. Since yearly fluctuations mostly dictate among-generation variation and daily fluctuations dictate within-generation variation, horizontal/vertical structures represent effects that are influenced by the frequency of the fluctuations, whereas diagonal structures in the heatmaps represent frequency-invariant effects. Results are averaged across the three beetle populations.

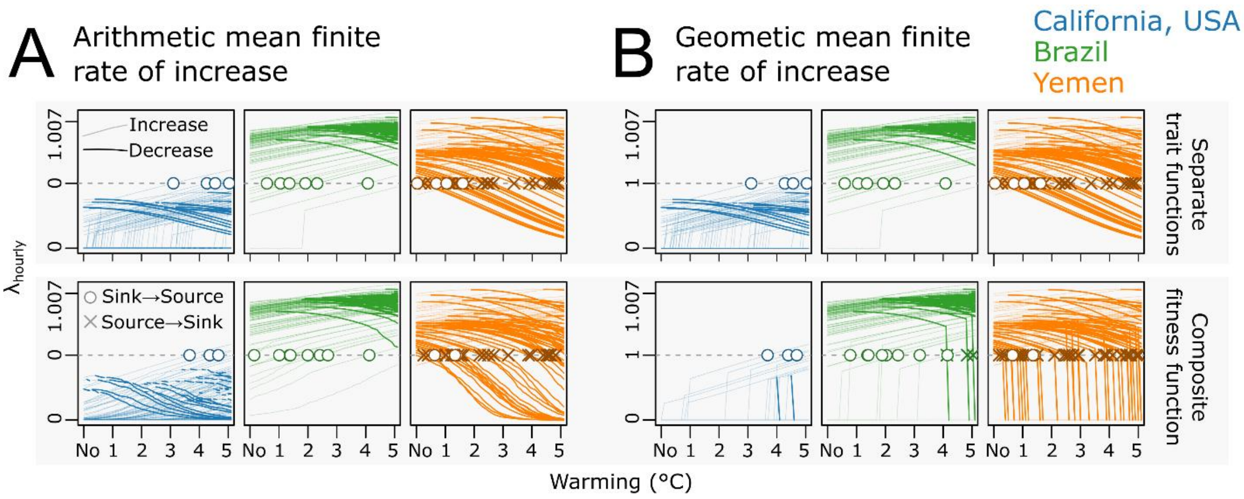


**Figure S5**. Differences among sites in hourly finite growth rates ($\lambda_{hourly}$) averaged over one year for different warming scenarios. (A) Arithmetic means. (B) Geometric means, with prevalent extinction events ($\lambda = 0$), particularly for the composite fitness function. Note the power scale of the y-axis.

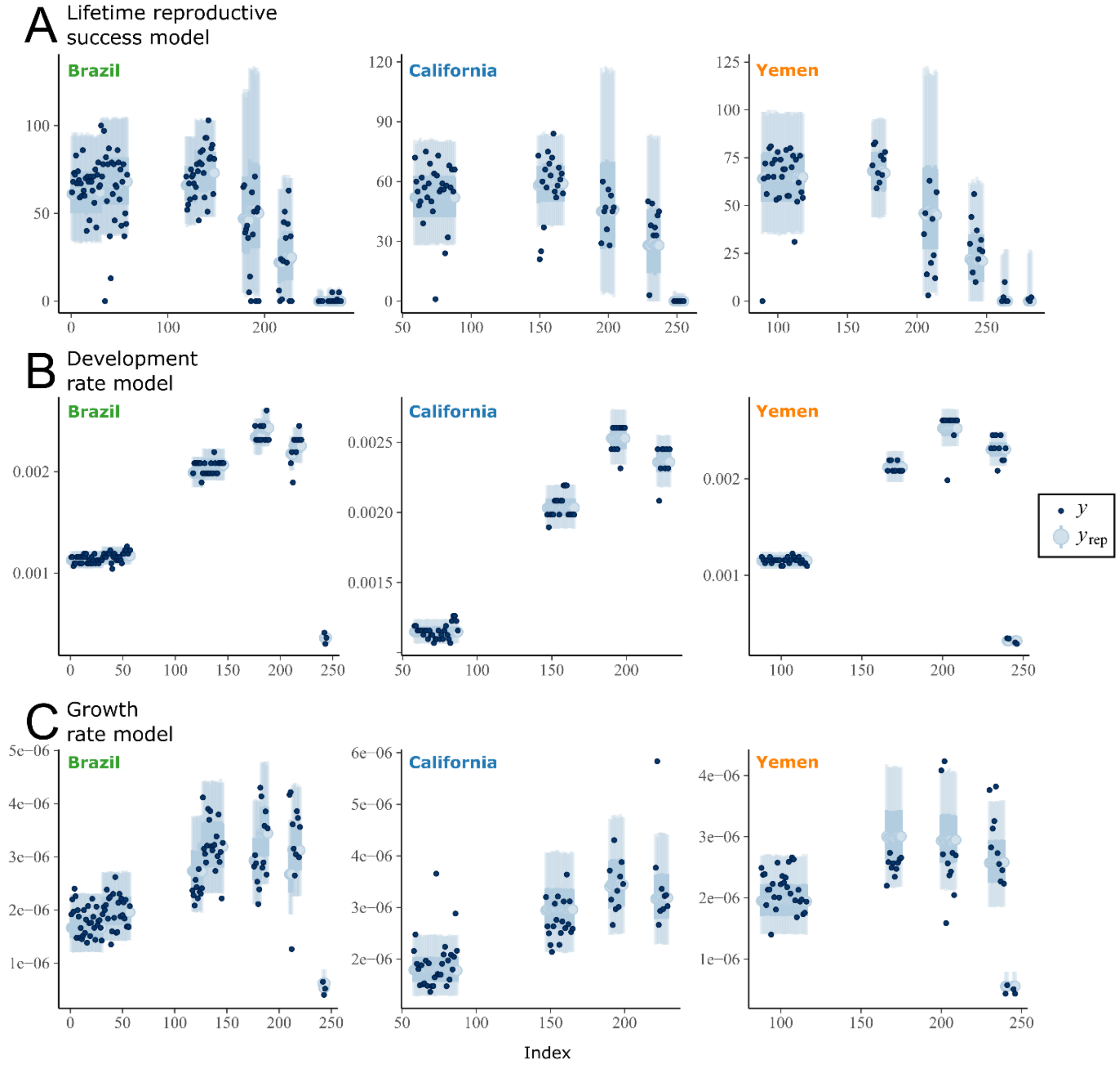


**Figure S6**. Comparisons between the distribution of observed data (dark points, y) and intervals representing distributions data simulated from the fitted models (blue bars, $y_{rep}$).

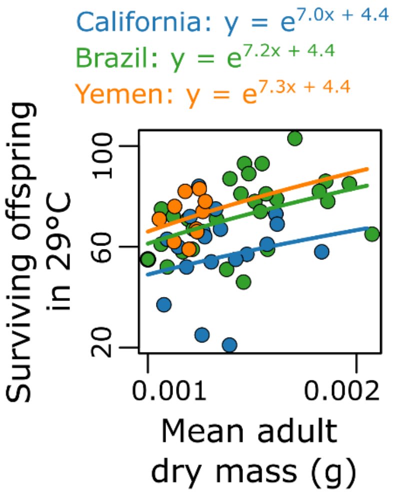


**Figure S7**. Scaling of the number of adult offspring (in 29°C) with average body mass. Points and lines are colored by origin: California, USA (blue), Brazil (green), and Yemen (orange).

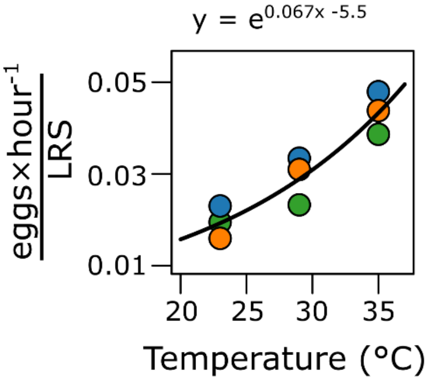


**Figure S8**. Relationship between temperature and the ratio of 1-hour oviposition rate to lifetime reproductive success (LRS).

**Table S1.** Bayesian posterior point estimates (modes), uncertainty (90% highest posterior density intervals), and priors for the parameters of the nonlinear models.

| | | Parameter | Mode | 90% HPI Lower | 90% HPI Upper | Prior |
|---|---|---|---|---|---|---|
| Lifetime reproductive success model | Viability | a | 4.32 | 2.79 | 6.88 | lognormal[mu = log(0.03), sigma = log(1.2)] |
| | | Topt | 0.0300 | 0.0209 | 0.0418 | gaussian[mu = 28, sigma = 2] |
| | | k | 28 | 25.6 | 29.7 | gaussian[mu = -4.5, sigma = 2] |
| | Fecundity | κ | -4.61 | -5.74 | -3.84 | lognormal[mu = log(2.5), sigma = log(1.5)] |
| | | $T_{opt}$ Brazil | 28.4 | 27.6 | 28.9 | gaussian[mu = 29, sigma = 5] |
| | | $T_{opt}$ California | 29.1 | 27.9 | 30.1 | gaussian[mu = 29, sigma = 5] |
| | | $T_{opt}$ Yemen | 28.1 | 26.5 | 28.8 | gaussian[mu = 29, sigma = 5] |
| | | $F_{max}$ Brazil | 70.6 | 66.3 | 76.0 | gaussian[mu = 70, sigma = 25] |
| | | $F_{max}$ California | 59.4 | 54.6 | 65.1 | gaussian[mu = 70, sigma = 25] |
| | | $F_{max}$ Yemen | 68.2 | 62.7 | 75.2 | gaussian[mu = 70, sigma = 25] |
| | | $T_b$ Brazil | 21.3 | 20.4 | 23.9 | lognormal[mu = log(22), sigma = log(1.1)] |
| | | $T_b$ California | 21.5 | 19.7 | 26.1 | lognormal[mu = log(22), sigma = log(1.1)] |
| | | $T_b$ Yemen | 22.1 | 20.8 | 25.7 | lognormal[mu = log(22), sigma = log(1.1)] |
| | | Brazil replicate effect (intercept)* | -0.0939 | -0.207 | 0.00512 | gaussian[mu = 1, sigma = 0] |
| Development rate model | | $T_{min}$ Brazil | 10.6 | 10.1 | 11.1 | gaussian[mu = 10, sigma = 3] |
| | | $T_{min}$ California | 8.89 | 7.00 | 10.8 | gaussian[mu = 10, sigma = 3] |
| | | $T_{min}$ Yemen | 11.1 | 10.7 | 11.5 | gaussian[mu = 10, sigma = 3] |
| | | $T_{opt}$ Brazil | 34.1 | 33.9 | 34.4 | gaussian[mu = 32.5, sigma = 3] |
| | | $T_{opt}$ California | 34.7 | 34.4 | 35.0 | gaussian[mu = 32.5, sigma = 3] |
| | | $T_{opt}$ Yemen | 34.0 | 33.8 | 34.3 | gaussian[mu = 32.5, sigma = 3] |
| | | $T_{max}$ Brazil | 42.9 | 41.8 | 44.2 | gaussian[mu = 37.5, sigma = 3] |
| | | $T_{max}$ California | 41.3 | 39.6 | 43.7 | gaussian[mu = 37.5, sigma = 3] |
| | | $T_{max}$ Yemen | 42.0 | 41.0 | 43.4 | gaussian[mu = 37.5, sigma = 3] |
| | | $r_{max}$ Brazil | 0.0024 | 0.00236 | 0.00244 | uniform[min = 0, max = 1] |
| | | $r_{max}$ California | 0.00253 | 0.00247 | 0.0026 | uniform[min = 0, max = 1] |
| | | $r_{max}$ Yemen | 0.00254 | 0.00248 | 0.0026 | uniform[min = 0, max = 1] |
| | | Brazil replicate effect (intercept)* | -0.0357 | -0.0539 | -0.0202 | gaussian[mu = 1, sigma = 0] |
| | | Residual st. dev. | 0.0446 | 0.041 | 0.0492 | scaled half student-t[d.f. = 3] † |
| Growth rate model | | $T_{min}$ Brazil | 11.1 | 9.56 | 12.3 | gaussian[mu = 10.6, sigma = 1]* |
| | | $T_{min}$ California | 7.94 | 6.18 | 9.73 | gaussian[mu = 8.89, sigma = 1]* |
| | | $T_{min}$ Yemen | 12.2 | 10.8 | 13.4 | gaussian[mu = 11.1, sigma = 1]* |
| | | $T_{opt}$ Brazil | 33.1 | 32.4 | 33.8 | gaussian[mu = 34.1, sigma = 1]* |
| | | $T_{opt}$ California | 34.1 | 33.3 | 35.0 | gaussian[mu = 34.7, sigma = 1]* |
| | | $T_{opt}$ Yemen | 31.8 | 31.2 | 32.6 | gaussian[mu = 34.0, sigma = 1]* |
| | | $T_{max}$ Brazil | 43.9 | 42.1 | 45.4 | gaussian[mu = 42.9, sigma = 1]* |
| | | $T_{max}$ California | 42.2 | 40.5 | 44.0 | gaussian[mu = 41.3, sigma = 1]* |
| | | $T_{max}$ Yemen | 43.3 | 41.9 | 45.1 | gaussian[mu = 42.0, sigma = 1]* |
| | | $r_{max}$ Brazil | $3.23\times10^{-6}$ | $3.07\times10^{-6}$ | $3.44\times10^{-6}$ | uniform[min = 0, max = 1] |
| | | $r_{max}$ California | $3.46\times10^{-6}$ | $3.24\times10^{-6}$ | $3.72\times10^{-6}$ | uniform[min = 0, max = 1] |
| | | $r_{max}$ Yemen | $3.15\times10^{-6}$ | $2.94\times10^{-6}$ | $3.36\times10^{-6}$ | uniform[min = 0, max = 1] |
| | | Brazil replicate effect (intercept)* | -0.16 | -0.234 | -0.0919 | gaussian[mu = 1, sigma = 0] |
| | | Residual st. dev. | 0.194 | 0.175 | 0.212 | scaled half student-t[d.f. = 3] † |

* Effect on the natural log scale   ** priors based on dev. rate posteriors   † *brms* default prior

**Table S2.** Point estimates (linear least squares), and associated uncertainty (95% confidence intervals) for the parameters of the linear models.

| Parameter | Estimate (LLS) | 95% CI lower | 95% CI upper |
|---|---|---|---|
| **Temperature-dependence of oviposition rate / LRS rate ratio** | | | |
| Intercept Brazil | 7.2 | 4.6 | 9.8 |
| Intercept California | 7.0 | 4.3 | 9.6 |
| Intercept Yemen | 7.3 | 4.6 | 9.9 |
| Slope | 0.44 | 0.047 | 0.84 |
| **Adult mass to number of offspring scaling** | | | |
| Intercept | -5.5 | -6.2 | -4.8 |
| Slope | 0.067 | 0.042 | 0.092 |
